# *Drosophila* HCN mediates gustatory homeostasis by preserving sensillar transepithelial potential in sweet environments

**DOI:** 10.1101/2024.02.06.579099

**Authors:** MinHyuk Lee, Se Hoon Park, Kyeung Min Joo, Jae Young Kwon, Kyung-Hoon Lee, KyeongJin Kang

## Abstract

Establishing transepithelial ion disparities is crucial for sensory functions in animals. In insect sensory organs called sensilla, a transepithelial potential, known as the sensillum potential (SP), arises through active ion transport across accessory cells, sensitizing receptor neurons such as mechanoreceptors and chemoreceptors. Because multiple receptor neurons are often co-housed in a sensillum and share SP, niche-prevalent overstimulation of single sensory neurons can compromise neighboring receptors by depleting SP. However, how such potential depletion is prevented to maintain sensory homeostasis remains unknown. Here, we find that the *Ih-*encoded hyperpolarization-activated cyclic nucleotide gated (HCN) channel bolsters the activity of bitter-sensing gustatory receptor neurons (bGRNs), albeit acting in sweet-sensing GRNs (sGRNs). For this task, HCN maintains SP despite prolonged sGRN stimulation induced by the diet mimicking their sweet feeding niche, such as overripe fruit. We present evidence that *Ih*-dependent demarcation of sGRN excitability is implemented to throttle SP consumption, which may have facilitated adaptation to a sweetness-dominated environment. Thus, HCN expressed in sGRNs serves as a key component of a simple yet versatile peripheral coding that regulates bitterness for optimal food intake in two contrasting ways: sweet-resilient preservation of bitter aversion and the previously reported sweet-dependent suppression of bitter taste.

## Introduction

Glia-like support cells exhibit close physical association with sensory receptor neurons, and conduct active transcellular ion transport, which is important for the operation of sensory systems^1^. In mammals, retinal pigment epithelial (RPE) cells have a polarized distribution of ion channels and transporters. They provide an ionic environment in the extracellular space apposing photoreceptors to aid their light sensing^2^. Likewise, knockdown of *Drosophila* genes encoding the Na^+^/K^+^ pump or a K^+^ channel in the supporting glial cells attenuates photoreceptors^3^. In addition to creating an optimal micro-environment, transepithelial potential differences (TEPs) are often generated to promote the functions of sensory organs. For example, the active K^+^ transport from the perilymph to the endolymph across support cells in the mammalian auditory system^4^ generates high driving forces that enhance the sensitivity of hair cells by increasing K^+^ and Ca^2+^ influx through force-gated channels. Similar designs have been found in insect mechanosensory^5,6^ and chemosensory organs^7,8^, providing models to study physiological principles and components of TEP function and regulation. Many insect sensory receptor neurons are housed in a cuticular sensory organ called the sensillum. Tight junctions between support cells separate the externally facing sensillar lymph from the internal body fluid known as hemolymph^9^. The active concentration of K^+^ in the dendritic sensillar lymph produces positive sensillum potentials (SP, +30∼40 mV) as TEPs, which are known to sensitize sensory reception in mechanosensation^10^ and chemosensation^11,12^.

Excitation of sensory neurons drains SP, accompanied by slow adaptation of the excited receptor neurons^6,12^. This suggests that immoderate activation of a single sensory neuron can deplete SP, which decreases the activities of neurons that utilize the potential for excitation. Each sensillum for mechanosensation and chemosensation houses multiple receptor neurons^1^. Therefore, overconsumption of SP by a single cell could affect the rest of the receptor neurons in the same sensillum, because the receptor neurons share the sensillum lymph. Indeed, the reduction of SP was proposed to have a negative effect on receptor neurons that are immersed in the same sensillar lymph; a dynamic lateral inhibition between olfactory receptor neurons (ORNs) occurs through “ephaptic interaction”, where SP consumption by activation of one neuron was proposed to result in hyperpolarization of an adjacent neuron, reducing its response to odorants^13,14^. As expected with this SP-centered model, ephaptic inhibition was reported to be mutual between *Drosophila* ORNs^13,15^, again because the ORNs are under the influence of a common extracellular fluid, the sensillar lymph. Such reciprocal cancellation between concomitantly excited ORNs may encode olfactory valence^16^ rather than lead to signal attenuation of two olfactory inputs. Furthermore, depending on neuron size, the lateral inhibition between ORNs can be asymmetric, albeit yet to be bilateral; larger ORNs are more inhibitory than smaller ones^13^. The size dependence was suggested to be due to the differential ability of ORNs to sink SP (referred to as local field potential in the study^13^), probably because larger cells have more membrane surface area and cell volume to move ions to or from the sensillar lymph.

Interestingly, gustatory ephaptic inhibition was recently found to be under a genetic, but not size-aided, regulation to promote sweetness-dependent suppression of bitterness^17^. This is accomplished by blocking one direction of ephaptic inhibition. The hyperpolarization-activated cation current in sGRNs through the *Ih-*encoded HCN is necessary to resist the inhibition of sGRNs laterally induced by bGRN activation. Furthermore, such unilateral ephaptic inhibition is achieved against cell size gradient^17^. Larger bGRNs are readily suppressed by the activation of smaller sGRNs, but not vice versa. Thus, HCN is implemented to inhibit bGRNs in terms of unilateral ephaptic inhibition when a bitter chemical is concomitantly presented with strong sweetness. Here, in addition to the ephaptic interaction, we find that the same HCN expressed in sGRNs promotes the activity of bGRNs as a means of homeostatic sensory adaptation, for which HCN prevents sGRNs from depleting SP even with the long-term exposure to the sweet-rich environment.

## Results

### HCN expressed in sweet-sensing GRNs is required for normal bitter GRN responses

The hair-like gustatory sensilla in the *Drosophila* labellum are categorized into L-, i-, and s-type based on their relative bristle lengths. Each sensillum contains 2 (i-type) or 4 (s- and L-type) GRNs along with a mechanosensory neuron. The i- and s-type bristle sensilla contain both an sGRN and a bGRN, while the L-type bristle sensilla contain an sGRN but no bGRN^18–21^. As a model of gustatory homeostasis, we mainly examined the i-type bristles using single sensillum extracellular recording^22–24^ because of their simple neuronal composition. Compared to WT (*w^1118^* in a Canton S background), we observed reduced spiking responses to 2 mM caffeine in two strong loss-of-function alleles of the HCN gene, *Ih^f03355^* ^25,26^ and *Ih^MI03196-TG4.0^/+* (*Ih- TG4.0/+*)^17,27^ (Fig.1A). Note that *Ih-TG4.0* is homozygous lethal^17^. A copy of the *Ih-*containing genomic fragment {*Ih*} rescued the spiking defect in *Ih^f03355^*. The GRN responses to 50 mM sucrose were not altered in *Ih* mutants (Fig.1B). Other bitter chemical compounds, berberine, lobeline, theophylline, and umbelliferone, also required *Ih* for normal bGRN responses (Fig. 1-figure supplement 1). Although we observe here that *Ih* pertains to bGRN excitability, *Ih* was previously found to be expressed in sGRNs but not bGRNs^17^. To test whether HCN expression in sGRNs is required for bGRN activity, GRN-specific RNAi knockdown of *Ih* was performed with either *Gr64f-*^28^ *or Gr89a-Gal4*^21^. *Ih* knockdown in sGRNs (*Gr64f-Gal4*), but not bGRNs (*Gr89a-Gal4*), led to reduced bGRN responses to caffeine (Fig.1C), indicating that HCN acts in sGRNs for a normal bGRN response. Unlike the results in *Ih* mutant alleles, the spiking response of *Ih*-knockdowned sGRNs (*Gr64f* cells) to 50 mM sucrose was increased (Fig.1D). To exclude the possibility that *Ih* is required for normal gustatory development, we temporally controlled *Ih* RNAi knockdown to occur only in adulthood, which produced similar results (Fig. 1-figure supplement 2). Such differential effects of gene disruptions and RNAi on sGRN activity will be discussed further below with additional results. Introduction of *Ih-RF* cDNA (Flybase id: FBtr0290109), which previously rescued *Ih* deficiency in other contexts^17,26^, to sGRNs but not bGRNs restored the decreased spiking response to 2 mM caffeine in *Ih^f03355^*, corroborating that sGRNs are required to express *Ih* for bGRN regulation (Fig.1E). Interestingly, ectopic cDNA expression in bGRNs of *Ih^f03355^* but not in sGRNs increased the spiking response to 50-mM sucrose compared to its controls (Fig.1F), although the same misexpression failed to raise the spiking to 2-mM caffeine. These results suggest not only that *Ih* innately expressed in sGRNs is necessary for the activity of bGRNs, but also that *Ih* expression in one GRN may promote the activity of the other adjacent GRN in *Ih-*deficient animals.

**Fig. 1.**
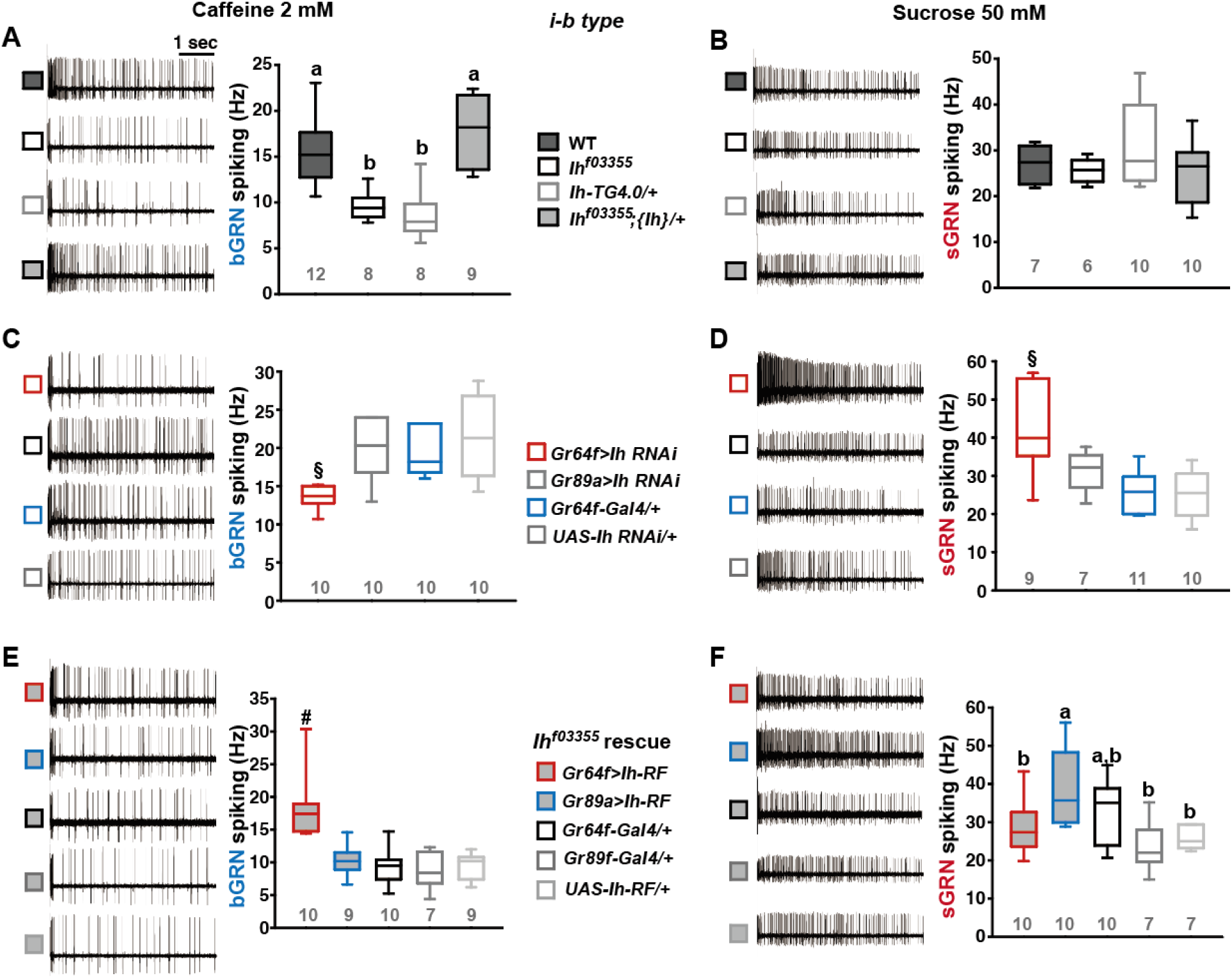
HCN is necessary for the normal activity of bitter-sensing GRNs (bGRNs), although expressed in sGRNs. Representative 5 sec-long traces of sensillum recording with either caffeine or sucrose at the indicated concentrations, shown along with box plots of spiking frequencies. (**A)** Caffeine-evoked bitter spiking responses of WT, the *Ih*-deficient mutants, *Ih^f03355^* and *Ih-TG4.0/+*, and the genomic rescue, *Ih^f03355^*;*{Ih}/+*. (**B**) Sucrose responses were similar among the genotypes tested in (**A**). (**C**) *Ih* RNAi knockdown in sGRNs, but not bGRNs, reduced the bGRN responses to 2-mM caffeine. (**D**) *Ih* RNAi knockdown in sGRNs increased the sGRN responses to 50-mM sucrose. (**E**) Introduction of the *Ih-RF* cDNA in sGRNs, but not bGRNs, of *Ih^f03355^* restored the bGRN response to 2-mM caffeine. (**F**) For sucrose responses, introduction of *Ih-RF* to bGRNs increased the spiking frequency. Letters indicate statistically distinct groups (a and b): Tukey’s test, p<0.05 (A), Dunn’s, p<0.05 (F). §: Welch’s ANOVA, Games-Howell test, p<0.05. #: Dunn’s test, p < 0.05. Numbers in gray indicate the number of tested naïve bristles, which are from at least 3 individuals.

### Loss of Ih in sGRNs reduced the sensillum potential in the gustatory bristle sensilla

We speculated that the *Ih-*dependent lateral boosting across GRNs might involve a functional link between GRNs. Such a physiological component could be sensillum potential (SP), since the sensillar lymph is shared by all GRNs in the sensillum and SP sets the spiking sensitivity^5^. SP is known as a transepithelial potential between the sensillum lymph and the hemolymph, generated by active ion transport through support cells (Fig. 2A, Left). To measure SP, we repurposed the Tasteprobe pre-amplifier to record potential changes in a direct current (DC) mode (see Methods for details), which was originally devised to register action potentials from sensory neurons. With the new setting, the contact of the recording electrode with a labellar bristle induced a rise in potential (Fig.2A, Right). The recording was stabilized within 20 sec, and a raw potential value was acquired as an average of the data between the time points, 20 and 60 sec after the initial contact (Fig.2A). After the examination of all the bristle sensilla of interest, the fly was impaled at the head to obtain the DC bias (also known as DC offset), which insects are known to exhibit in the body independent of SP^29^ (Fig.2B). To examine whether the DC bias varies at different body sites, we surveyed the DC bias at four different locations of individual animals, the abdomen, thorax, eye, and head. This effort resulted in largely invariable DC bias readings (Fig.2B,C). Next, the sensillum potential was obtained by subtracting the DC bias from the raw potential value (Fig.2A). We also found that we could reduce the apparent SP by deflecting the bristle sensillum by ∼45° (Fig.2D-F), activating the sensillum’s mechanosensory neuron. When we performed the same experiment with *nompC^f00642^*, a loss-of-function allele of *nompC* that encodes a mechanosensory TRPN channel^30^, this reduction in SP disappeared (Fig.2D-F).

**Fig. 2.**
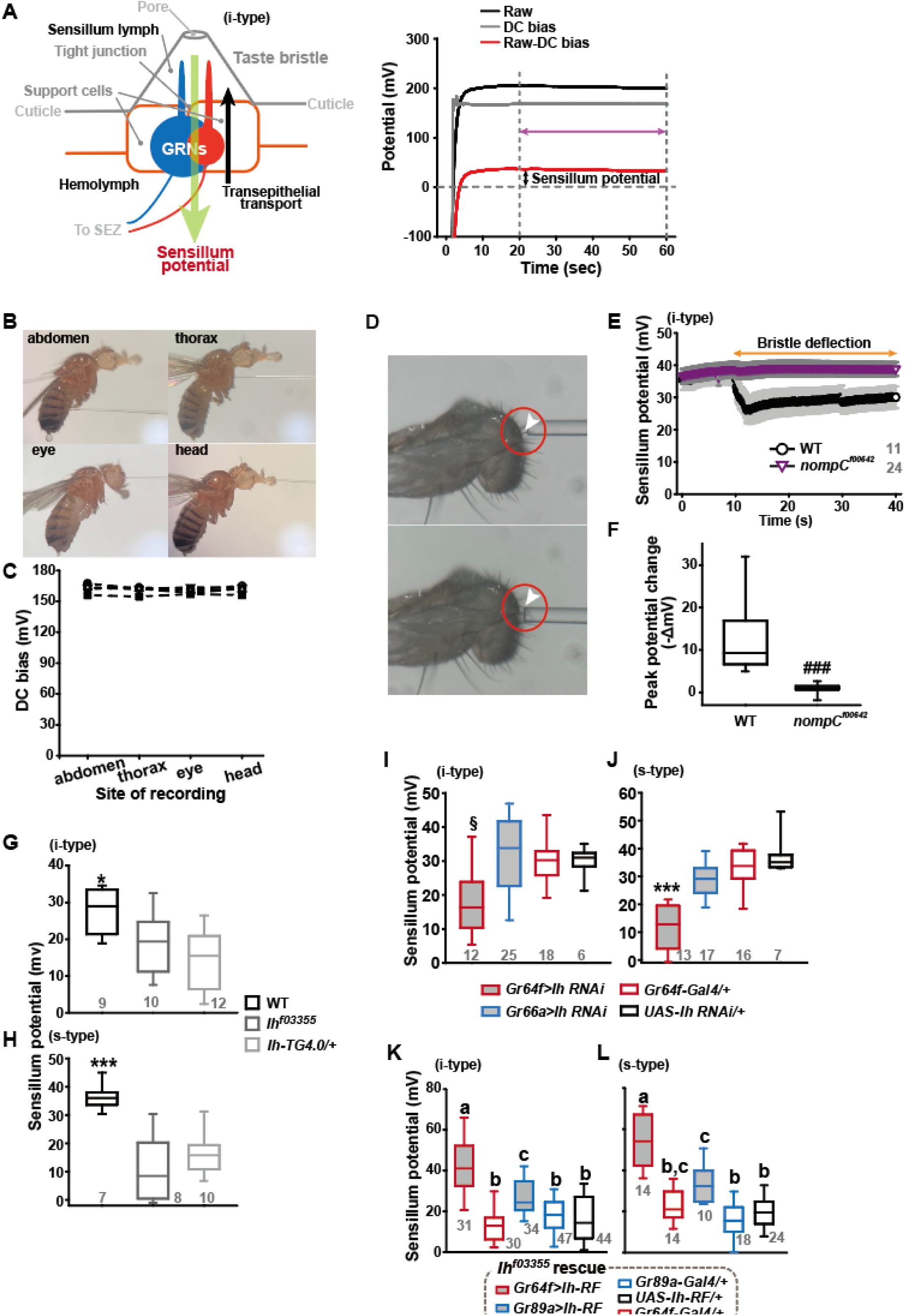
Sensillum potential (SP) is reduced in HCN-deficient animals. (**A**) Schematic diagram illustrating the sensillum potential in the taste bristle sensilla (Left). Black upward arrow indicates ion transport by pumps and transporters in support cells from the hemolymph to the sensillum lymph, which are physiologically separated by tight junctions between support cells. The resulting transcellular disparity of ions leads to a positive sensillum potential. Representative traces of potentials measured to evaluate SP (Right). Raw: the potential reading upon the contact of the recording electrode with the sensillum bristle tip (black). DC bias: the potential reading upon impalement of the head by the recording electrode (gray). Red line indicates the difference between raw and DC bias, which represents the sensillum potential. The values resulting from the subtraction of the data between 20 to 60 sec after the initial contact (time indicated by the purple double headed arrow) were averaged to determine SP. (**B**) Photographs of impaled flies for DC bias determination at indicated sites. (**C**) DC bias values obtained from indicated body parts. There is no statistical significance between the body sites (ANOVA Repeated Measures). (**D**) Photos before (top) and after (bottom) deflection of an i-type bristle. (**E**) Sensillum potential traces as a function of time from WT and *nompC^f00642^*. Bristle bending started at 10 sec, and the duration is marked by an orange double headed arrow. (**F**) The peak SP changes of WT and *nompC^f00642^*were compared. (**G** and **H**) SP was reduced in i- (G) and s-type (H) bristles of the indicated *Ih*-deficient mutants, relative to WT. (**I** and **J**) *Ih* RNAi in sGRNs reduced SPs of the i- and s-type bristles. (**K** and **L**) The SP of *Ih^f03355^* was restored by *Ih-RF* expression in GRNs (red for sGRNs, blue for bGRNs). ###: Dunn’s, p<0.001. * and ***: Tukey’s, p<0.05 and p<0.001, respectively. §: Welch’s ANOVA, Games-Howell test, p<0.05. Letters indicate statistically distinct groups: Tukey’s test, p < 0.05. Numbers in gray indicate the number of naive bristles tested in at least three animals.

Suggesting the role of *Ih* in SP regulation, *Ih^f03355^* (∼19 and ∼10 mV for i-type and s-type sensilla, respectively) and *Ih-TG4.0/+* (∼15 and ∼16 mV for i-type and s-type sensilla, respectively) exhibited reduced mean SPs compared to WT in the i-type (Fig.2G) and s-type (Fig.2H) bristle sensilla (∼28 mV and ∼36 mV, respectively). We also examined whether the SP reduction could be attributed to the lack of *Ih* in sGRNs through GRN-specific *Ih* RNAi knockdown. This revealed that *Ih* is necessary in sGRNs for the sensilla to exhibit normal SP levels (Fig.2I,J). The SP reduction observed in both bristle types of *Ih^f03355^* could be fully restored by expressing the *Ih-RF* cDNA in sGRNs (*Gr64f-Gal4* cells). Mean SPs were measured to be ∼42 and ∼54 mV in i-type and s-type bristles, respectively (Fig.2K,L). Interestingly, ectopic expression of the cDNA in bGRNs by *Gr89a-Gal4* also significantly rescued the SP defect of *Ih^f03355^* to the level of mean SPs (∼27 and ∼33 mV in i-type and s-type bristles, respectively) comparable to those in WT. The greater extent of SP defect restoration in *Ih^f03355^* by *Ih-RF* expressed in sGRNs than bGRNs indicates that *Ih-RF* is more effective at upholding SP in sGRNs than in bGRNs under our experimental conditions. Furthermore, the successful rescue by *Ih-RF* in bGRNs also shows that *Ih* can regulate SP in any GRN (Fig.2K,L).

### Inactivation of sGRNs raised both bGRN activity and SP, which was reversed by Ih deficiency

Since it is in sGRNs that HCN regulates the bGRN responsiveness to caffeine, we suspected that the activity of sGRNs may be closely associated with the maintenance of bGRN excitability. In line with this possibility, the *Gr64af* deletion mutant, which lacks the entire *Gr64* gene locus and is severely impaired in sucrose and glucose sensing^31–33^, showed increased bGRN responses to various bitters in labellar gustatory bristle sensilla compared to WT (Fig.3A). Furthermore, silencing sGRNs (*Gr5a-Gal4* cells) by expressing the inwardly rectifying potassium channel, Kir2.1^34^, phenocopied *Gr64af* in response to 2-mM caffeine stimulating the i-type bristles (Fig.3B). This increased responsiveness of bGRNs is unlikely due to positive feedback resulting from the sGRN inactivation through the neural circuitry in the brain, because the tetanus toxin light chain (TNT) expressed in sGRNs, which blocks chemical synaptic transmission^35^, failed to raise bGRN activity (Fig.3C). Strikingly, when we combined the sGRN-hindering genotypes (*Gr5a>Kir2.1* and *Gr64af*) with the *Ih* alleles *Ih^f03355^* or *Ih-TG4.0,* we found that the sGRN inhibition-induced increase in bGRN activity in response to caffeine could be commonly relieved by the disruptions in the *Ih* gene (Fig.3B,E). This result suggests that HCN suppresses sGRN activation, while HCN expressed in sGRNs is required for unimpaired bGRN activity (Fig.1C,E). Interestingly, Kir2.1-induced inactivation of sGRNs (*Gr64f-Gal4* cells) dramatically increased the mean SP of the i-type bristles to ∼53 mV, compared to ∼29 and ∼35 mV of *Gal4* and *UAS* controls, respectively (Fig.3D), and the impairment of sucrose-sensing in the *Gr64af* mutants also resulted in increases of mean SPs (Fig.3F, ∼56 and ∼53 mV in the i- and s-bristles of *Gr64af*, compared to ∼30 and ∼36 mV of WT, respectively). Thus, inactivating sGRNs in two different ways increased SP in the i- and s-type gustatory bristles, similar to the effect on bGRN activity described earlier. Such repeated parallel shifts of bGRN activity and SP were again obtained in the combined genotypes between *Gr64af* and *Ih^f03355^*or *Ih-TG4.0/+* (Fig.3F); the SP increased in *Gr64af* descended to WT levels when combined with *Ih^f03355^*and *Ih-TG4.0/+*, similar to what occurred with bGRN activity in *Gr64af* (Fig.3E). These results suggest that *Ih* gene expression suppresses sGRNs, upholding both bGRN activity and SP, similar to the genetic alterations that reduce sGRN activity.

**Fig. 3.**
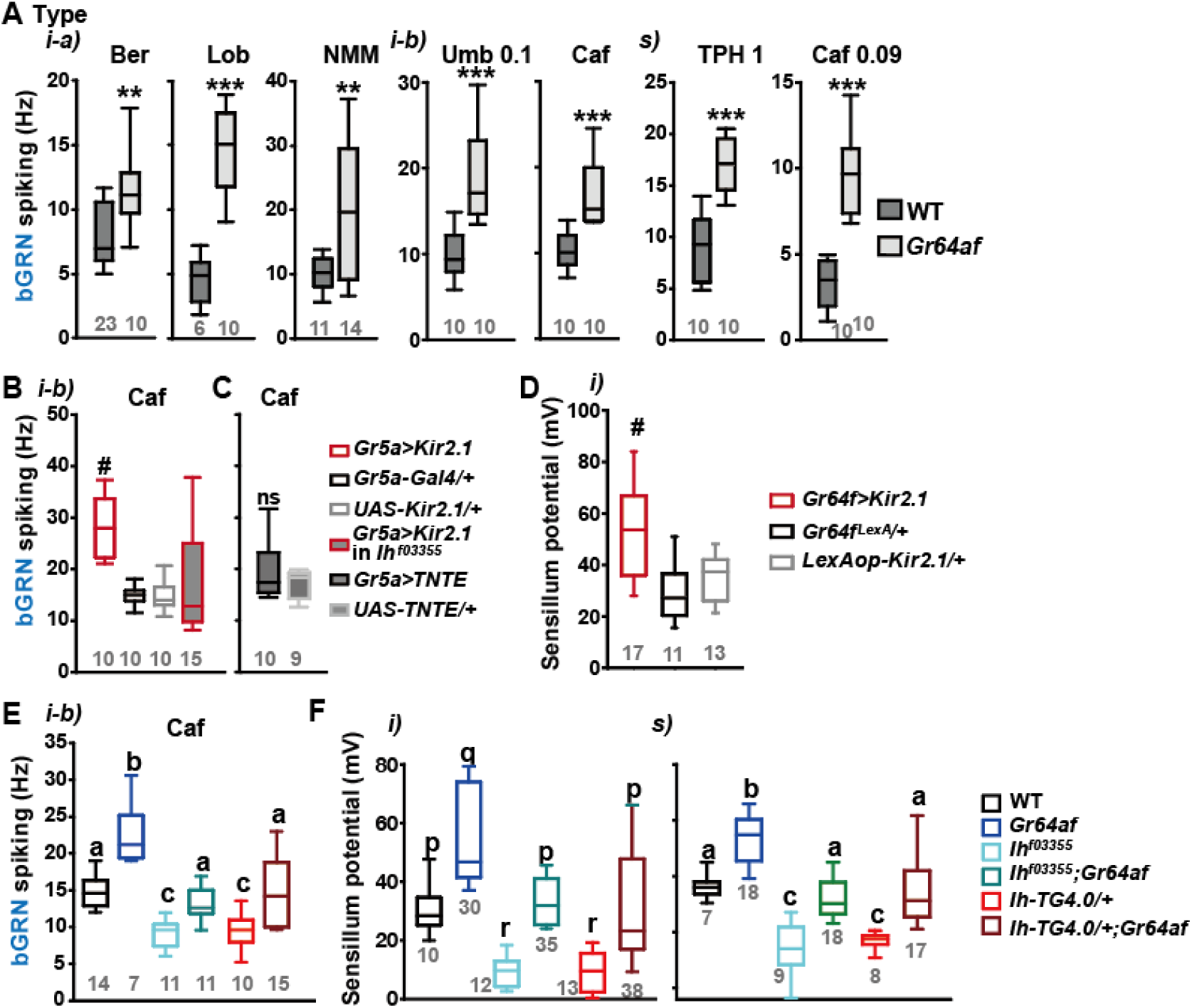
Inactivation of sGRNs raises bGRN activity and SP, both of which are reversed by *Ih* deficiency. (**A**) The bGRN spiking was increased in response to the indicated bitters in *Gr64af* mutants impaired in sucrose and glucose sensing. Ber: 0.5, Lob: 0.5, NMM: 2, Caf: 2 (i-type) and 0.09 (s-type), Umb: 0.1, TPH: 1 mM. * and ***: Student’s t-test, p<0.01 and p<0.001, respectively. (**B** and **C**) Silencing by Kir2.1 (B), but not blocking chemical synaptic transmission (C), in sGRNs increased the spiking of bGRNs stimulated by 2-mM caffeine, which was reversed in *Ih^f03355^* (B). #: Dunn’s, p<0.05. (**D**) Silencing sGRNs by Kir2.1 increased SP. #: Dunn’s, p<0.05. (**E**) The increased bGRN spiking in *Gr64af* was restored to WT levels by *Ih* deficiencies. Letters indicate significantly different groups (Tukey’s, p<0.05). Caffeine 2 mM was used (B, C and E). (**F**) Regardless of bristle type, SP was increased upon sGRN inactivation, which was reduced by *Ih* deficiencies. **p**-**r**: Dunn’s test, p<0.05. **a-c**: Welch’s ANOVA, Games-Howell test, p<0.05. Numbers in gray indicate the number of naïve bristles in at least 3 animals.

Water GRNs are co-housed with sGRNs in L-type bristles in the labellum, responding to hypo-osmolarity with the aid of *ppk28* and promoting water drinking^36,37^. We tested whether *Ih*- dependent SP regulation occurs in these bristles to maintain the sensitivity of water GRNs by using a low concentration of the electrolyte tricholine citrate (TCC) at 0.1 mM. Interestingly, L-type bristles of *Ih^f03355^* showed reduced spike frequencies in response to this hypo-osmolar electrolyte solution compared to WT (Fig. 3-figure supplement 1A). This reduction was restored in the genetic rescue line. Additionally, SP in these bristles was increased in *Gr64af* but decreased in the two *Ih* alleles, and the combination of the *Gr64* and *Ih* mutations restored SP to the level of WT (Fig. 3-figure supplement 1B), as observed with other sensillar bristles above. Finally, *Ih-RF* restored SP in *Ih^f03355^* when expressed in sGRNs but not bGRNs, as expected from the absence of bGRNs in L types (Fig. 3-figure supplement 1C). Thus, *Ih-*dependent SP regulation is universal in all bristle sensilla of the labellum and likely important for the function of GRNs neighboring sGRNs.

### HCN delimits excitability of HCN-expressing GRNs, and increases SP

By misexpressing *Ih-RF* in bGRNs of WT flies, we investigated how HCN physiologically controls HCN-expressing GRNs (Fig.4A). The genetic controls, *Gr89a-Gal4/+* and *UAS-Ih-RF/+*, exhibited mutually similar dose dependencies saturated at 2- and 10-mM caffeine, revealing the maximal caffeine responses at these concentrations. Interestingly, the ectopic expression reduced bGRN activity at these high caffeine concentrations (Fig.4A). The flattened dose dependence suggests that ectopically expressed HCN suppresses strong excitation of bGRNs. In contrast, sGRNs were upregulated by the misexpression of *Ih* in bGRNs with increased spiking in response to 10- and 50-mM sucrose (Fig.4B), implying that *Ih* increases the activity of the neighboring GRN by reducing that of *Ih*-expressing GRNs. On the other hand, the *Ih-RF*-overexpressing sGRNs in *Gr64f-Gal4* cells significantly decreased only the response 5 sec after contacting 50-mM sucrose (Fig.4C, the second 5-sec bin, Fig. 4-figure supplement 1), probably because of native HCN preoccupying WT sGRNs. Although bGRNs were repressed by misexpressing *Ih-RF*, the mean SPs increased to ∼40 and ∼37 mV in the i- and s-type bristles, respectively, compared to controls with mean SPs of 22-25 mV (Fig.4D). These results from misexpression experiments corroborate the postulation that sGRNs are suppressed by expressing HCN. To confirm that sGRNs are suppressed by native HCN, the impact of GRN-specific *Ih* RNAi knockdown on sGRNs was quantitatively evaluated (Fig.4E). *Ih* RNAi in sGRNs (*Gr64f*-*Gal4* cells) led to increased mean spiking frequencies by ∼10 Hz in response to 1-, 5-, and 10- mM as well as 50-mM sucrose compared to *Ih* RNAi in bGRNs (*Gr66a-Gal4* cells) and genetic controls, highlighting the extent to which HCN natively expressed in sGRNs suppresses sGRN excitability. In contrast, SP, necessary for GRN sensitization, was observed above to be reduced by *Ih* RNAi in sGRNs but not bGRNs (Fig.2I,J). Thus, these datasuggest that HCN innately reduces the spiking frequencies of sGRNs even at relatively low sucrose concentrations, 1, and 5 mM. This is similar to the suppressive effect of *Ih-RF* misexpressed in bGRNs at relatively high caffeine concentrations, but differs in that the misexpression did not alter bGRN activity in response to low caffeine concentrations, 0.02 and 0.2 mM (Fig.4A), implying a complex cell-specific regulation of GRN excitability.

**Fig. 4.**
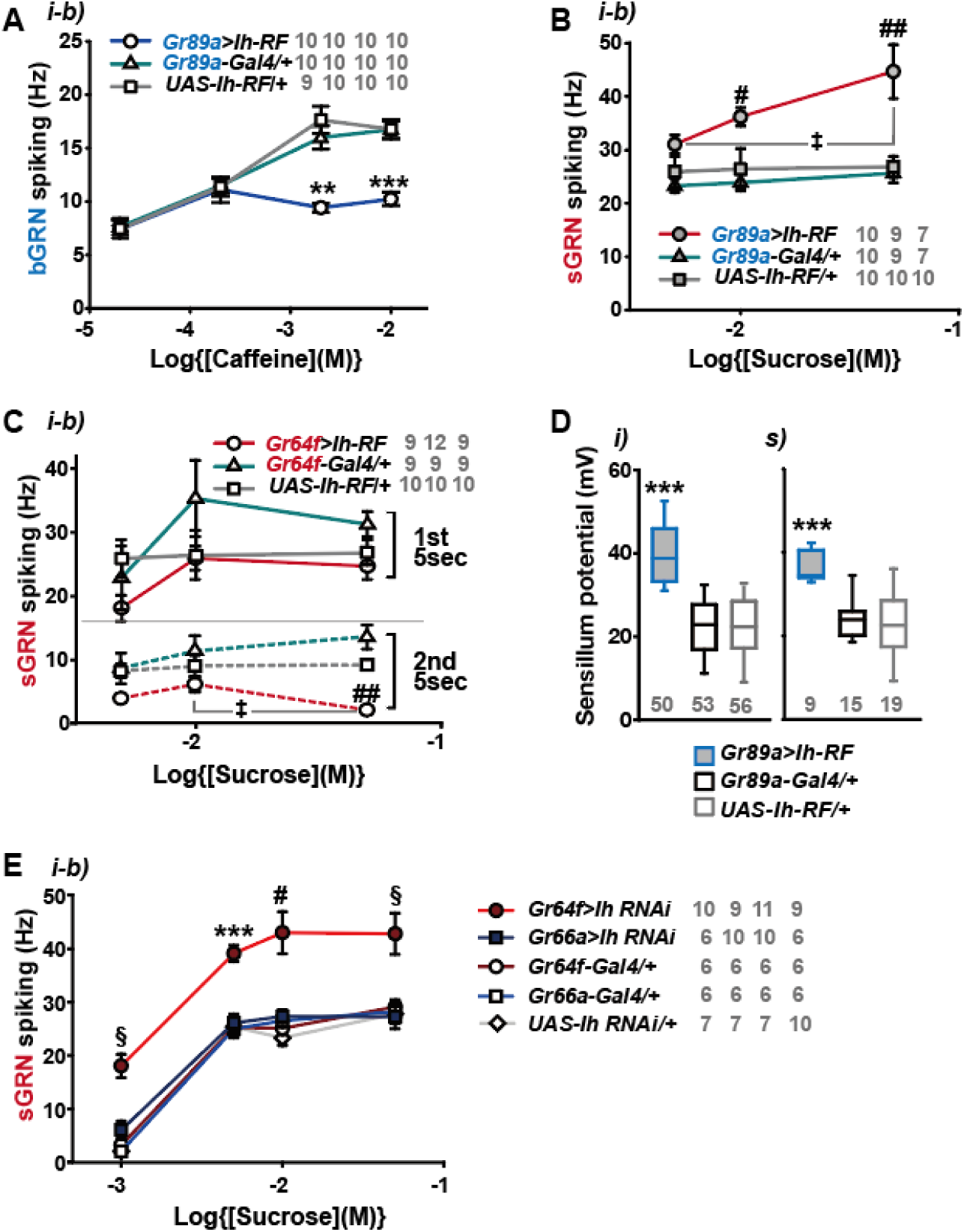
HCN suppresses HCN-expressing GRNs and increases SP. (**A**) HCN misexpressed in bGRNs flattened the dose dependence to caffeine. (**B**) HCN ectopically expressed in bGRNs elevates sGRN responses to sucrose. (**C**) Overexpression of HCN in sGRNs reduced the sGRN responses to sucrose 5 sec after the initial contact. (**D**) *Ih* misexpression in bGRNs increased SP in i- and s-type bristles, which correlates with laterally increased sGRN activity (B). (**E**) *Ih* RNAi knockdown in sGRNs (*Gr64f-Gal4* cells) dramatically elevates spiking frequencies in response to 1-, 5-, 10-, and 50-mM sucrose. *, **, and ***: Tukey’s, p<0.05, p<0.01, and p<0.001, respectively (A,D,E). # and ##: Dunn’s, p<0.05 and p < 0.01 between genotypes, respectively (B,C,E). ‡: Dunn’s, p < 0.05 between responses to different sucrose concentrations (B,C). §: Welch’s ANOVA, Games-Howell test, p<0.05 (E). The numbers in grey indicate the number of tested naïve bristles in at least three animals.

### Sweetness in the food leads to reduction of SP, bGRN activity, and bitter avoidance in Ih-deficient animals

Typically, we performed extracellular recordings on flies 4-5 days after eclosion, during which they were kept in a vial with fresh regular cornmeal food containing ∼400 mM D-glucose. The presence of sweetness in the food would impose strong and frequent stimulation of sGRNs for extended period, potentially requiring the delimitation of sGRN excitability for the homeostatic maintenance of gustatory functions. To investigate this possibility, we fed WT and *Ih^f03355^* flies overnight with either non-sweet sorbitol alone (200 mM) or a sweet mixture of sorbitol (200 mM) + sucrose (100 mM). Although sorbitol is not sweet, it is a digestible sugar that provides *Drosophila* with calories^38^. We found that the sweet sucrose medium significantly reduced caffeine-induced bGRN responses in both genotypes compared to the sorbitol only medium, but *Ih^f03355^* bGRN spike frequencies were decreased to the level significantly lower than WT (Fig.5A), as seen above with the cornmeal food (Figs.1A,C and 3E). This suggests that the reduced bGRN activity in the mutants may result from prolonged sGRN excitation. The SP reduction was induced by 1-hr incubation with the sweet sucrose medium in both WT and *Ih^f03355^* to a similar degree. However, the *Ih* mutant showed a more severe depletion of SPs compared to WT after 4 hrs of sweet exposure (Fig.5B) as observed with the cornmeal food (Figs.2 and 3F). Even on the sorbitol food, the SP in *Ih^f03355^* was significantly decreased compared to WT. This may be attributed to the loss of HCN, which is known to stabilize the resting membrane potential^39^. Following overnight sweet exposure, SPs of WT and *Ih^f03355^* were recovered to similar levels after 1-hr incubation with sorbitol only food. However, it was after 4 hrs on the sorbitol food that the two lines exhibited SP levels similar to those achieved by overnight incubation with sorbitol only food (Fig. 5B). These results indicate that SP depletion by sweetness is a slow process, and that the dysregulated reduction and recovery of SPs in *Ih^f03355^*manifest only after long-term conditioning with and without sweetness, respectively.

**Fig. 5.**
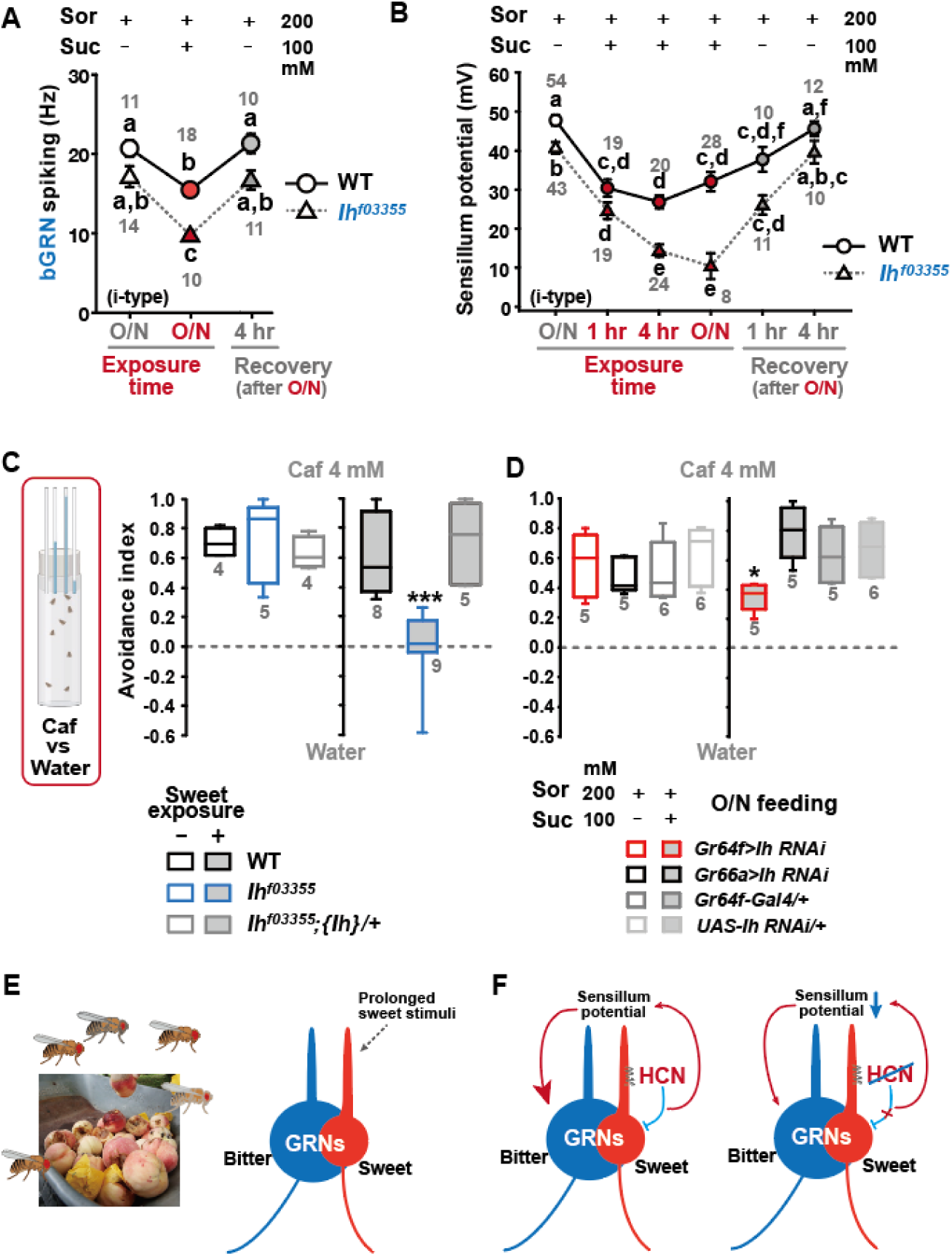
Sweetness in the diet decreases SP, bGRN activity, and bitter avoidance. (**A**) Sweetness in the media reduced the 2-mM caffeine-evoked bGRN spiking, which was fully recovered in four-hour incubation with sorbitol only food. *Ih^f03355^* was affected by the type of the media more severely than WT. O/N: overnight incubation with sorbitol only (grey) or sucrose food (red). (**B**) The SP of *Ih^f03355^* bristle sensilla showed dysregulated reduction after four-hour and overnight incubation on sweet media. These reductions started to be recovered in one-hour feeding and were nearly fully recovered in four-hour feeding on the indicated sorbitol only food. (**C**) Caffeine (Caf) avoidance was assessed with capillary feeder assay (CAFE). *Ih* is required for robust caffeine avoidance for flies maintained on sweet cornmeal food (sweet exposure +: filled boxes). *Ih^f03355^* flies avoided 4 mM caffeine like WT flies when separated from sweet food for 20 hrs (blank boxes). (**D**) *Ih* RNAi knockdown in sGRNs (*Gr64f-Gal4*) but not bGRNs (*Gr66a-Gal4*) led to relatively poor avoidance to caffeine after feeding on the sweet diet with sucrose. Suc: sucrose, and Sor: sorbitol Letters indicate statistically distinct groups: a-f, Dunn’s, p < 0.05 (A,B). * and ***: Tukey’s, p<0.05 and <0.001, respectively. (E) Illustration depicting the flies’ sweet feeding niche in overripe fruit (Left), leading to prolonged exposure of sGRNs to the sweetness (Right). (F) A schematic model of gustatory homeostasis in *Drosophila* bristle sensilla. Despite the prolonged sweetness in the environment robustly and frequently stimulating sGRNs, the sGRN activity is moderated by HCN to preserve the sensillum potential, which is required for normal bGRN responsiveness (Left). When HCN in sGRNs is incapacitated, sGRNs can become overly excited by sweetness of overripe fruit and deplete the sensillum potential, resulting in decreased bGRN activity and bitter avoidance (Right).

To assess the behavioral implications of HCN-assisted preservation of SP and bGRN activity, flies were exposed long-term to sweetness on a regular sweet cornmeal diet (sweet exposure-positive), and then subjected to a CAFE with an 8-hr choice between water and 4 mM caffeine solution. Note that sucrose was not used in CAFE, because the presence of sweet stimuli was shown to suppress bGRNs^17^. Indicative of reduced bitter sensitivity, *Ih^f03355^* flies showed dramatically decreased caffeine avoidance, relative to WT (Fig.5C). In contrast, when flies were removed from the cornmeal food for 20 hrs, both WT and *Ih^f03355^* showed similarly robust bitter avoidance. The defect observed in the *Ih* mutant on the sweet cornmeal diet could be rescued by reintroducing a genomic fragment covering the *Ih* locus ({*Ih*}). These results were recapitulated with other bitters, lobeline and theophylline (Fig. 5-figure supplement 1). To examine whether caffeine avoidance requires *Ih* expression in sGRNs, CAFE was performed with GRN-specific RNAi knockdown of *Ih*. For the RNAi experiments, flies were kept overnight on either the non-sweet diet with sorbitol (200 mM) or the sweet diet with additional sucrose (100 mM). *Ih* knockdown in sGRNs, but not bGRNs, led to a deficit in the avoidance only when the flies were on the sweet diet, indicating that HCN expression in sGRNs is necessary for robust caffeine avoidance in a sweet environment (Fig.5D). Therefore, the sweetness of the diet can compromise the function of bGRNs co-housed with sGRNs in the same sensilla, which is mitigated by HCN expression in sGRNs. Such a role of HCN is essential for bitter avoidance of flies, considering their likely prolonged exposure to sweetness in their natural habitat of overripe fruit (Fig. 5E).

## Discussion

Our results provide multiple lines of evidence that HCN suppresses HCN-expressing GRNs, thereby sustaining the activity of neighboring GRNs within the same sensilla (Fig. 5F). We propose that this modulation occurs by restricting SP consumption through HCN-dependent neuronal suppression rather than via chemical and electrical synaptic transmission. The lack of increased bGRN activity with TNT expression in sGRNs, coupled with the increase observed with Kir2.1 expression (Fig.3B,C), indicates minimal involvement of synaptic vesicle-dependent transmission. The possibility of a neuropeptide-dependent mechanism is unlikely, given our ectopic gain-of-function studies (Fig. 4). To explain the misexpression results with neuropeptide pathways, both s- and bGRNs must be equipped with the same set of a neuropeptide/receptor system, which is incompatible with the inverse relationship between the two GRNs in excitability observed in Fig. 1C,D, Fig. 1-figure supplement 2B, and Fig. 3. Furthermore, this inverse relationship argues against electrical synapses through gap junctions, which typically synchronize the excitability of pre- and postsynaptic neurons. Therefore, our findings propose an unconventional mechanism of neuronal interaction.

HCNs are encoded by four different genes in mammals^39,40^, and are known to be present in mammalian sensory receptor cells. In cochlear hair cells, HCN1 and HCN2 were reported to form a complex with a stereociliary tip-link protein^41^, while in vestibular hair cells, HCN1 is essential for normal balance^42^. HCN1 was also immunostained in cone and rod photoreceptors, as well as retinal bipolar, amacrine, and ganglion neurons, with deletion of the encoding gene resulting in prolonged light responses^43^. A subset of mouse taste cells was labeled for HCN1 and HCN4 transcripts and proteins^44^, similar to our observation of selective HCN expression in *Drosophila* GRNs. HCN2 is expressed in small nociceptive neurons that mediate diabetic pain^45^. However, the precise roles of HCNs in regulating these respective sensory physiologies remain to be elucidated.

HCN is well-known for its ‘funny’ electrophysiological characteristics, stabilizing the membrane potential^39,40^. As a population of HCN channels remains open at the resting membrane potential, HCN serves to suppress neuronal excitation in two ways. First, it increases the inward current required to depolarize the membrane and trigger action potentials, owing to the low membrane input resistance resulting from the HCN-dependent passive conductance. Second, the closing of HCN induced by membrane depolarization counteracts the depolarization, since the reduction of the standing cation influx through HCN is hyperpolarizing. Conversely, HCNs also allow neurons to resist membrane hyperpolarization because the hyperpolarization activates HCNs to conduct depolarizing inward currents. Consequently, HCN channels effectively dampen fluctuations in membrane potential, whether they lead to depolarization or hyperpolarization. Our findings in this study align with the former property of HCNs, as *Drosophila* HCN is essential for moderating sGRN excitation to preserve SP and bGRN activity when flies inhabit in sweet environments. On the other hand, our previous study showed that HCN-dependent resilience to hyperpolarizing inhibition of sGRNs lateralizes gustatory ephaptic inhibition to dynamically repress bGRNs, when exposed to strong sweetness together with bitterness^17^. Thus, depending on the given feeding contexts, the electrophysiological properties of HCN in sGRNs lead to playing dual roles with opposing effects in regulating bGRNs. The stabilization of membrane potential by HCNs was reported to decrease the spontaneous activity of neurons, as evidenced by miniature postsynaptic currents suppressed by presynaptic HCNs^46^. In this regard, the lower SP observed in the *Ih^f03355^* labellar bristles than that of WT, even on the nonsweet sorbitol food (Fig.5B), may be attributed to the more facile fluctuations in resting membrane potential which could regulate the consumption of SP (further discussion below).

Cell-specific knockdown of *Ih* in sGRNs led to increased sGRN responses to 50-mM sucrose (Fig.1D), although disruptions of the *Ih* locus did not (Fig.1B). This inconsistency may stem from differences between alleles and the RNAi knockdown in residual *Ih* expression or in *Ih-*deficient sites. The lack of *Ih* in sGRNs can induce two different effects in neuronal excitation: 1) easier depolarization of sGRNs due to the loss of standing HCN currents at rest (as suggested in Fig.4E) and 2) a decrease of receptor-mediated inward currents, expected due to SP reductions (Fig.2G-L). Assuming that some level of HCN expression may persist in RNAi knockdowns compared to mutants, these opposing effects on sGRN excitability may largely offset each other in response to 50-mM sucrose in the *Ih* mutants, but not in the knockdowned flies. The ectopic introduction of *Ih-RF* into bGRNs of *Ih^f03355^* significantly increased the mean SP compared to the control genotypes (Fig.2K,L), leaving sGRNs devoid of functional *Ih*. This genotype allows the examination of sGRNs lacking *Ih*, with SP unimpaired, which is supposed to reflect the net effect of *Ih* on sGRN excitability excluding the influence from reduced SP. Interestingly, the ectopic rescue resulted in elevated firing responses to 50-mM sucrose compared to the cDNA rescue in sGRNs (Fig.1F), a proper control with *Ih* expression and SP both unimpaired. On the other hand, the differing sites of *Ih* deficiency might create the inconsistency. The protein trap reporter *Ih-TG4.0-Gal4* previously showed widespread expression of HCN in the labellum, including non-neuronal cells, implying the possibility of unknown bGRN-regulating HCN-dependent mechanisms, potentially harbored in nonneuronal cells. Overall, our cell-specific loss-of-function and gain-of function studies advocate that HCN suppresses HCN-expressing GRNs, which thereby increases SP to promote the activity of the neighboring GRNs.

Only the dendrites of GRNs face the sensillar lymph, separated from the hemolymph by tight junctions between support cells^9^. The inward current through the ion channels that respond to sensory reception in the dendrites is thought to be a major sink for SP^5,12^, consistent with the incremented SP in the *Gr64af* mutant lacking the sucrose-sensing molecular receptor^17,31^. Based on these points, it was somewhat unexpected that the membrane potential regulator HCN preserved SP, yet implying that the sensory signaling in the dendrite is likely under voltage-dependent control. In line with HCN, shifting the membrane potential toward the K^+^ equilibrium by overexpressing Kir2.1 in sGRNs upregulated bGRN activity and SP (Fig.3), corroborating that the membrane potential in sGRNs is a regulator of the sensory signaling cascade in the dendrites. Note that the sensillum lymph contains high [K^+^]^5,7^, which would not allow strong inactivation of sGRNs and SP increases if Kir2.1 operates mostly in the dendrites. The increases in SP, coinciding with the apparent silencing of sGRNs by Kir2.1^17^, propose that lowering the membrane potential in the soma and the axon suppresses the consumption of SP probably by inhibiting the gustatory signaling-associated inward currents in the dendrite. Para, the *Drosophila* voltage-gated sodium channel, was reported to be localized in the dendrites of mechanosensitive receptor neurons in *Drosophila* chordotonal organs^47^. Similarly, *Drosophila* voltage-gated calcium channels have been studied in dendrites^48–50^, implying that membrane potential may be an important contributor to the sensory signaling in dendrites.

There are ∼14,500 hair cells in the human cochlea at birth^51^. These hair cells share the endolymph in the scala media (cochlear duct), representing a case of TEP shared by a large group of sensory receptor cells. Since HCNs were found to be unnecessary for mechanotransduction itself in the inner ear^52^, they may play a regulatory role in fine-tuning the balance between the endocochlear potential maintenance and mechanotransduction sensitivity for hearing, as in the *Drosophila* gustatory system. Multiple mechanosensory neurons are found to be co-housed also in *Drosophila* mechanosensory organs such as hair plates and chordotonal organs^5^. Given that each mechanosensory neuron is specifically tuned to detect different mechanical stimuli such as the angle, velocity, and acceleration of joint movement^53^, some elements of these movements may occur more frequently and persistently than others in a specific ecological niche. Such biased stimulation would require HCN-dependent moderation to preserve the sensitivity of other mechanoreceptors sharing the sensillar lymph. We showed that ectopic expression of *Ih* in bGRNs also upheld SP and the activity of the neighboring sGRNs, underscoring the independent capability of HCN in SP preservation. Despite such an option available, the preference for sGRNs over bGRNs in HCN-mediated taste homeostasis implies that *Drosophila melanogaster* may have ecologically adapted to the high sweetness^54^ prevalent in their feeding niche, such as overripe or fermented fruits^55^. It would be interesting to investigate whether and how respective niches of various insect species differentiate the HCN expression pattern in sensory receptor neurons for ecological adaptation.

In this report, we introduce a peripheral coding design for feeding decisions that relies on HCN. HCN operating in sGRNs allows uninterrupted bitter avoidance, even when flies reside in sweet environments. This is achieved in parallel with an ephaptic mechanism of taste interaction by the same HCN in sGRNs, whereby bitter aversion can be dynamically attenuated in the simultaneous presence of sweetness^17^. Further studies are warranted to uncover similar principles of HCN-dependent adaptation in other sensory contexts. It would also be interesting to explore whether the role of HCN in the sensory adaptation consistently correlates with lateralized ephaptic inhibition between sensory receptors, given that sensory cells expressing HCN can resist both depolarization and hyperpolarization of the membrane.

## Acknowledgments

We would like to thank Dr. Kyuhyung Kim for helpful comments, and Drs. Amrein, H, Scott, K, Moon SJ, and KDRC/BDRC stock/resource centers for sharing fly lines as indicated in Methods and Materials.

## Author contributions

Conceptualization: MHL, KJK

Methodology: MHL, KJK, SHP, JYK

Investigation: MHL, KJK, KMJ, SHP, KL

Visualization: MHL, KJK, SHP, KL

Funding acquisition: KJK,

Project administration: JYK, KMJ, KL

Supervision: KJK, JYK, KMJ

Writing – original draft: MHL, KJK

Writing – review & editing: JYK, KMJ, SHP

## Data and materials availability

All data are available in the main text or the supplementary materials.

