## Supplementary material for "*Drosophila* HCN mediates gustatory homeostasis by preserving sensillar transepithelial potential in sweet environments": method and supp figures

Materials and Methods

Fly strains

The *w^1118^* line in a Canton-S background was used as wild type. *Gr64f-Gal4* was provided by Dr. Hubert Amrein, and *Gr5a-Gal4* by Dr. Kristin Scott. *Gr64af* is a gift from Dr. Seok Jun Moon. *UAS-Ih RNAi* (#58089)*,* a duplicate of the *Ih* locus (denoted as *{Ih}* in the main text, #89744), *Ih^f03355^* (#85660) and *Ih-TG4.0* (#76162) were acquired from Bloomington Drosophila Stock Center (#stock number). The *UAS-Ih-RF* line was previously generated by Korea Drosophila Resource Center (<http://kdrc.kr>) by site-specific recombination into attP49b (3R), for which we cloned *Ih* cDNA through reverse transcription^1^. The genotypes used in this study are linked to Fig. panels in the table below.

Extracellular recordings

In vivo extracellular recordings were performed by the tip-dip method as detailed previously^2,3^. Each of the i-a, i-b, s-b type sensillum of 3-5 day-old flies were identified from the sensillum map described elsewhere^4^. The reference electrode was filled with HL3.1 solution^5^. The recording electrode contained tastants solubilized in the electrolyte 2 (i-type) or 30 (L- and s-type) mM tricholine citrate (TCC). The concentrations of bitter chemicals were indicated in the corresponding figure legend. The spiking frequency (Hz) was calculated from the number of spikes in the first 5 second or the second 5 second as indicated, and compared between genotypes or experimental conditions. The signals picked up by the electrodes were amplified by the preamplifier Tasteprobe (Syntech) and digitized at a rate of 20 kb/sec by PowerLab with Labchart software (ADInstruments). The number of experiments indicated in figures are the number of naïve bristles tested. The naïve bristles were from at least three different animals.

Sensillum potential recordings

Media with or without sweetness were prepared as follows; the sorbitol medium consisted of 0.5% agarose and 200 mM sorbitol, while the sweet medium contained 0.5% agarose, 200 mM sorbitol and 100 mM sucrose. Flies were kept overnight on these media before the experiment. For SP recordings, the recording electrode contained 2 mM TCC as the electrolyte, and Tasteprobe was set to record in “pass-through” mode with DC (in the High-Pass filter window) and 100-ms zeroing time settings. Amplified signals were digitized at a rate of 100 Hz using PowerLab/Labchart. First, differential potentials were measured between a recording electrode on a taste sensillum and a reference electrode inserted into the labellum as performed for the extracellular bristle sensillum recordings. DC bias^6^ was measured by impaling the recording electrode into the thorax of the same animals used for SP measurements. DC bias was subsequently subtracted from the initial readouts of the differential potential to evaluate SP (Fig. 2A). The resulting SPs were averaged during 40-sec long recording 20 sec after initial contact, which were subsequently used for further analyses.

Bitter avoidance assay

Twenty flies, aged 3-5 days and consisting of 10 males and 10 females, were used to assess bitter avoidance using capillary feeder assay (CAFE). To test the bitter sensitivity of each genotype of interest in feeding behavior, flies were kept on regular cornmeal food or starved on nonsweet water-soaked Kimwipes overnight, and then given a choice between water and 4 mM caffeine for 8 hours. For RNAi experiments, 200-mM sorbitol is used in nonsweet food and sweet food, the latter of which included 100-mM sucrose in addition. Avoidance indices were obtained as the net volume fraction of water consumption subtracted by the volume fraction of caffeine ingestion.

Statistics

Statistical calculation was performed using Sigmaplot 14.5 (Systat Software). The sample sizes and the statistical tests are indicated in each figure or in the legend. Normal distribution and heteroskedasticity were assessed using Shapiro-Wilk and Brown-Forsythe tests, respectively, before parametric tests. When these tests were failed, non-parametric tests were performed. However, for some cases, heteroskedasticity with normality led us to perform Welch’s t-test (Sigmaplot 14) or Welch’s ANOVA. The latter was followed by Games-Howell test as a parametric analysis using the Excel spreadsheet available at [www.biostathandbook.com/welchanova.xls](http://www.biostathandbook.com/welchanova.xls). No outlier was excluded for statistical analyses.

| REAGENT | SOURCE | IDENTIFIER |
| --- | --- | --- |
| Experimental models: Organisms/strains | | |
| Cantonized *w^1118^* |  |  |
| *Gr64af* | Dr. Moon at Yonsei U. |  |
| *Ih^f03355^* | Bloomington Drosophila Stock Center | BDSC: 85660; Flybase: FBti0051182 |
| *Mi{Trojan-GAL4.0}IhMI03196-TG4.0 (*indicated as *Ih-TG4.0* in the text*)* | Bloomington Drosophila Stock Center | BDSC: 76162; Flybase: FBti0187533 |
| Duplicate of *Dp(2;3)GV-CH321-22I11 (*indicated as *{Ih})* | Bloomington Drosophila Stock Center | BDSC: 89744; Flybase: FBab0048672 |
| *Gr5a-Gal4* | Dr. Scott at UC Berkeley |  |
| *Gr64f-Gal4* | Dr. Amrein at TAMU |  |
| *Gr89a-Gal4* | Dr. Carlson at Yale |  |
| *UAS-Kir2.1* | Bloomington Drosophila Stock Center | BDSC: 6495 |
| *UAS-TNTE* | Bloomington Drosophila Stock Center | BDSC: 28837 |
| *UAS-Ih-RF* | This study |  |
| *UAS*-*Ih* RNAi | Bloomington Drosophila Stock Center | BDSC: 58089; Flybase: FBst0058089 |
| *nompC^f00642^* | Korea Drosophila Resource Center | KDRC: K3137; Flybase: FBti0041920 |
| Software and algorithms |  |  |
| LabChart 8 | AD Instrument | <https://www.adinstruments.com> |
| SigmaPlot 14.0 | Systat Software Inc | https://systatsoftware.com/ |

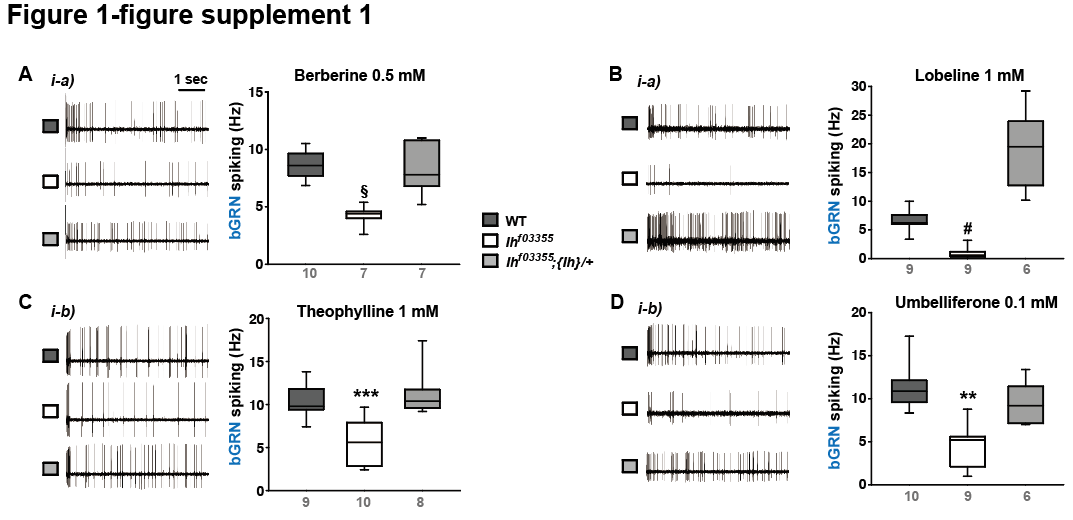

**Fig. 1-figure supplement 1.** ***Ih* is required for spiking responses to various bitter chemical compounds.** Representative 5 sec-long traces of sensillum recording in WT, *Ih^f03355^* and a genomic rescue are shown along with box plots of spiking frequencies for indicated bitters, such as berberine (A), lobeline (B), theophylline (C), and umbelliferone (D). §: Welch’s ANOVA, Games-Howell test, p<0.05. #: Dunn’s test, p < 0.05. ** and ***: Tukey’s, p<0.01 and p<0.001, respectively. Numbers in gray indicate the number of naïve bristles tested in at least three animals.

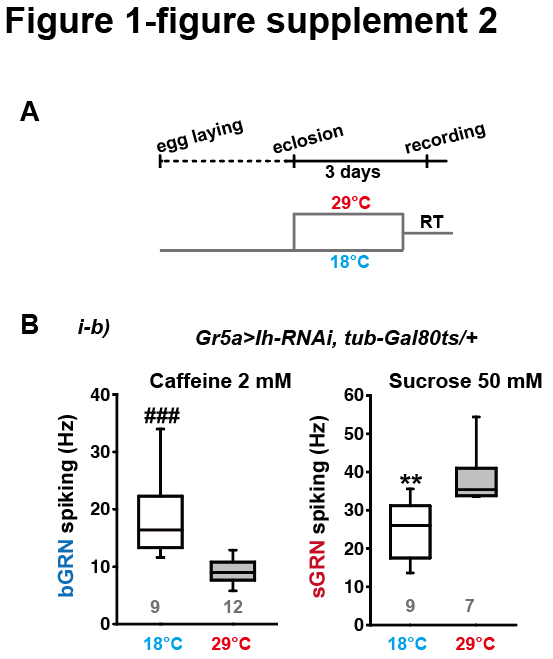

**Fig. 1-figure supplement 2. *Ih RNAi* knockdown in adulthood reduces spiking frequencies in response to 2 mM caffeine but increases spiking frequencies to 50 mM sucrose.** (A) Schematic diagram depicting the design of temporal control of the RNAi. (B) Box plots of spiking frequencies obtained with indicated bitter and sweet chemical compounds at temperatures permissive and non-permissive for Gal80ts. ###: Dunn’s, p<0.001. **: Tukey’s, p<0.01. Numbers in gray indicate the number of naïve bristles tested in at least three animals.

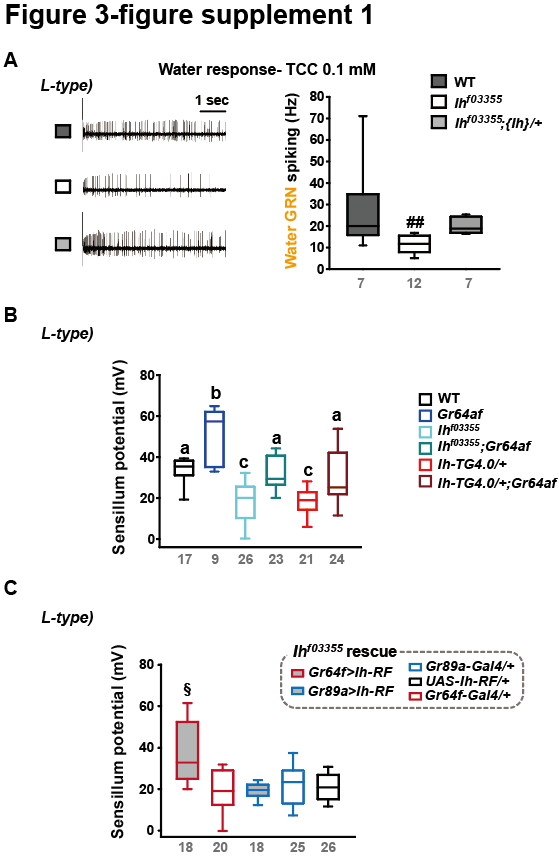

**Fig. 3-figure supplement 1. Water GRNs rely on the sensillum potential (SP) guarded by HCN in the L-type bristles.** (A) Water GRN activity evoked by 0.1 mM tricholine citrate (TCC) was appraised in WT, *Ih^f03355^* and a genomic rescue. The representative traces (Left) and box plots of spiking frequencies (Right) are shown. ##: Dunn’s, p<0.01. (B) SP in L-type bristles is reduced in *Ih-*deficient mutants but increased in *Gr64af*. Combination of *Ih* and *Gr64af* deficiencies cancels the respective effects, moving SPs towards the level observed in WT, as shown in i- and s-type bristles. Letters, a to c, indicate statistically distinct groups: Tukey’s, p<0.05. (C) Introduction of the *Ih-RF* cDNA in sGRNs, but not in bGRNs, restored SP in *Ih^f03355^*. §: Welch’s ANOVA, Games-Howell test, p<0.05. Numbers in gray indicate the number of naïve bristles tested in at least three animals.

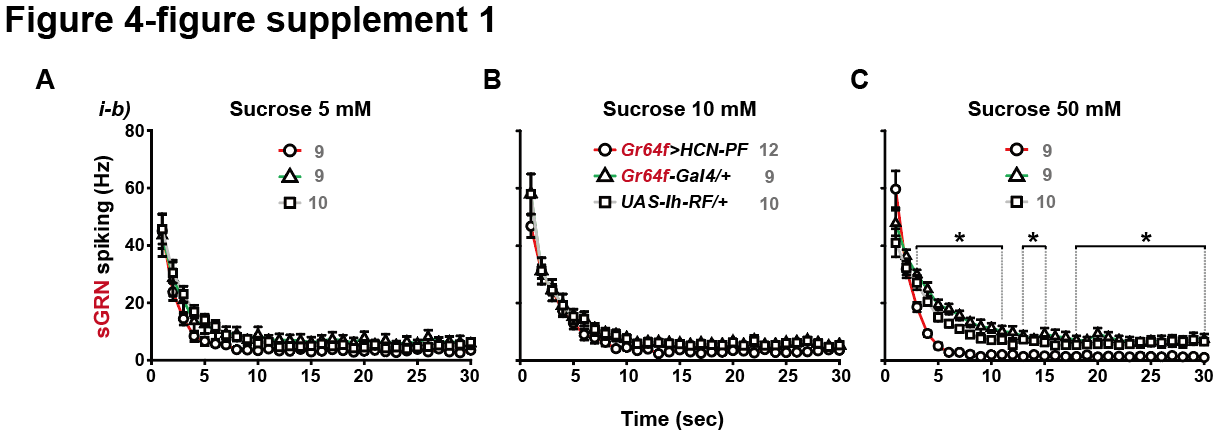

**Fig. 4-figure supplement 1. Overexpression of *Ih-RF* in WT sGRNs suppresses their spiking responses to 50 mM sucrose in a delayed manner.** Post-stimulus spiking frequencies binned every second are shown for sucrose concentrations, 5, 10 and 50 mM (A, B and C, respectively). *: p<0.05, Tukey’s, Dunn’s or Games-Howell test, depending on the data distribution and variance. The data from i-b type bristles of *Gr64f>Ih-RF* are significantly different from the genetic controls within the three indicated ranges. Numbers in gray indicate the number of naïve bristles tested in at least three animals. See Fig. 4C for a different style of data presentation.

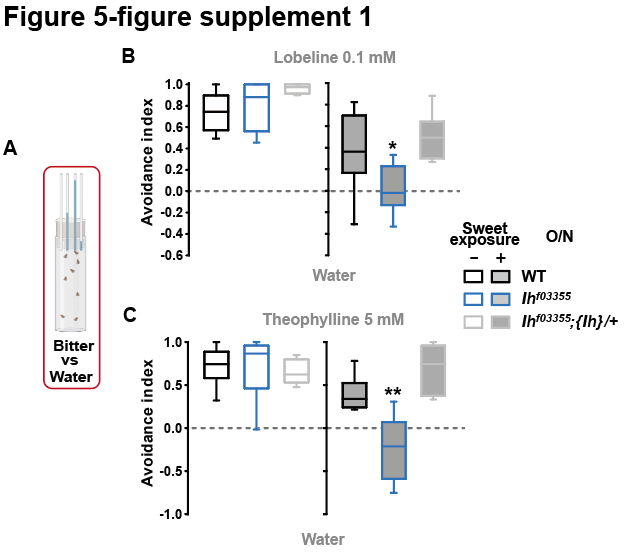

**Fig. 5-figure supplement 1. Feeding avoidance to lobeline and theophylline is reduced in *Ih^f03355^* following prior exposure to sweetness.** (A) Bitter avoidance was evaluated by CAFE. (B and C) *Ih* is required for avoidance to indicated bitters for flies maintained on sweet cornmeal food­ (sweet exposure +: filled boxes) but not for flies separated from sweetness for 20 hours (sweet exposure -: blank boxes). * and **: Tukey’s, p<0.05 and 0.01, respectively. Numbers in gray indicate the number of naïve bristles tested in at least three animals.
